# Dysfunction of telomeric Cdc13-Stn1-Ten1 simultaneously activates the DNA damage and spindle checkpoints

**DOI:** 10.1101/2024.08.06.606774

**Authors:** Nathalie Grandin, Michel Charbonneau

## Abstract

Telomeres, the ends of eukaryotic linear chromosomes, are composed of repeated DNA sequences and specialized proteins, with the conserved Cdc13/CTC1-Stn1-Ten1 (CST) telomeric comple providing chromosome stability via telomere end-protection and regulation of telomerase accessibility. In the present study, *SIZ1*, coding for a SUMO E3 ligase, and *TOP2* (Top2 is a SUMO target for Siz1 and Siz2) were isolated as extragenic suppressors of temperature-sensitive mutants of *Saccharomyces cerevisiae* CST. *ten1*-*sz, stn1*-*sz* and *cdc13*-*sz* mutants were next isolated on the basis of being sensitive to intracellular Siz1 dosage. In parallel, strong negative genetic interactions between mutants of CST and septins were identified, septins being noticeably sumoylated through action of Siz1. The temperature-sensitive arrest in these new mutants of CST was dependent on the G2/M Mad2-mediated and Bub2-mediated spindle checkpoints as well as on the G2/M Mec1-mediated DNA damage checkpoint. Our data suggest the existence of yet unknown functions of the telomeric Cdc13-Stn1-Ten1 complex related with mitotic spindle positioning and/or spindle assembly that could be further elucidated by studying these new *ten1*-*sz, stn1*-*sz* and *cdc13*-*sz* mutants.

## 1. Introduction

Telomeres are composed of TG-rich DNA sequences and specialized binding proteins, forming together high order assemblies. Telomeres protect the ends of the linear chromosomes from being recognized as DNA double strand breaks by the DNA damage signaling and DNA repair machineries [1]. In dividing cells, telomeres shorten at each cell division due to incomplete lagging-strand DNA replication, oxidative damage and exonucleolytic processes [2]. Excessive telomere erosion elicits DNA damage that activates cell cycle checkpoints, leading to senescence or apoptosis. The action of telomerase, a specialized reverse transcriptase [3, 4], at telomere ends or the activation of the alternative lengthening of telomeres pathway (ALT) [5] compensate for this erosion of telomeres - notably in cancer cells. The existence of numerous telomere-impacting genetic disorders implicated in cancer, aging and other diseases [6-8] have contributed to the importance of the field of telomere biology research during the last three decades.

In vertebrates, telomere protection is provided mainly by shelterin, a complex of six proteins (TRF1, TRF2, POT1, TIN2, TPP1 and RAP1) that prevents inappropriate recombination and fusion between telomeres. Shelterin subunits also have complementary roles in telomere replication and length regulation [9-11]. A similar complex exists in the fission yeast *Schizosaccharomyces pombe* [12], while a somewhat simpler protection complex, consisting mainly of the Cdc13, Stn1 and Ten1 proteins (the CST complex) is present in the budding yeast *Saccharomyces cerevisiae* [13, 14]. The *S. cerevisiae* CST complex is a telomere-specific Replication Protein A-like complex that plays a central role in telomere homeostasis and chromosomal end protection through its telomere capping and telomerase activation and inhibition functions [15-19]. Strikingly, orthologs of the three *S. cerevisiae* CST proteins are found in humans and mouse, as well as in the plant *Arabidopsis thaliana* [20, 21]. In *S. pombe*, Stn1 and Ten1, but not Cdc13/CTC1, have been identified as CST subunits [22].

Human and mouse CST were initially discovered by virtue of homology of their STN1 subunit to *S. cerevisiae* Stn1, followed by spectrometry analyses to identify the CTC1 and TEN1 subunits [20]. Human CST limits telomerase access at telomeres [23] and associates with shieldin at damaged telomeres, regulating in association with Polα, the fill-in of the resected overhangs to facilitate DNA repair [24]. Importantly, hCST subunits were found by immunofluorescence to localize at only a fraction of telomeres and the hypersensitivity of mutants of hCST to DNA damaging agents indicated a function in ensuring genome-wide replication restart in response to DNA replication stress and replication fork stalling [20, 25-27]. Finally, hCST has a role in the initiation of DNA replication in association with MCM proteins [28] and also physically associates with the cohesin complex, perhaps to prevent premature cohesion loss between chromosomes at stalled replication forks [29]. Recently too, *A. thaliana* TEN1 was shown to act as a heat-shock-induced molecular chaperone, potentially protecting CTC1 from forming aggregates [30].

Given the conservation of CST during evolution, a complete understanding of its functions is even more needed than before [10, 11, 27, 31]. In *S. cerevisiae* CST, Cdc13 has the most important role due to the high specificity of its single-stranded DNA binding capacity, ensuring telomere replication by recruiting telomerase and telomere end protection by associating with Stn1 [15, 32, 33]. Stn1 is essential for telomere end protection and also functions in telomere length regulation [17], being responsible, in association with DNA Polα, for the fill-in of the strand previously elongated by telomerase [34, 35]. Ten1, however, has no known specific telomeric function besides being, like Cdc13 and Stn1, essential for telomere end protection and telomere length regulation [18]. It also acts to improve Cdc13 stability at the telomeres by direct binding [36]. In addition, *S. cerevisisae* Cdc13-Stn1-Ten1 might have another (extratelomeric) function, in regulating transcription elongation [37], a function that has recently been proposed to be defective in human colorectal cancers [38].

Here we set out to develop a novel strategy to uncover novel loss-of-function mutants of CST. To do this, a temperature-sensitive synthetic *CDC13*-*TEN1*-*STN1* fusion gene, called *CST1*, was expressed in a strain in which the three corresponding (essential) genes had been deleted. Genetic suppressor screens allowed us to isolate *SIZ1*, coding for a SUMO E3 ligase, and *TOP2*, coding for topoisomerase II, as extragenic suppressors of these mutants. In parallel, we identified strong negative genetic interactions between mutants of CST and septins. Noticeably, septins are sumoylated through the action of Siz1. Septins are GTP-binding, cytoskeletal proteins, conserved from yeast to humans, that were first identified in budding yeast for their essential role in cytokinesis and play also critical roles in diverse functions including morphogenesis, mitosis, cell migration, ciliogenesis and exocytosis [39, 40]. Septins form heterooligomeric complexes polymerizing end-to-end into filaments that can be further organized into higher order structures, such as rings and hourglasses, depending on cell types or cell cycle stages [41, 42] [. Moreover, septins act as scaffolds to recruit cytokinesis factors to the site of cell division and regulate constriction of the contractile actomyosin ring [43]. In budding yeast, septins were the first substrates reported to undergo sumoylation [44, 45] and in humans but not in yeast, sumoylation of septins is critical for septin filament bundling and cytokinesis [40].

In addition, our data show that the damage generated by these novel mutants of CST is sensed by the two major G2/M spindle checkpoints, as well as by the major G2/M DNA damage checkpoint. The present study suggests the existence of (a) novel function(s) for the telomeric CST complex, which might be to maintain genome integrity in anaphase through activation of Siz1 and modulation of cell cycle progression by the septins.

## 2. Materials and methods

### 2.1. Yeast strains and media

*Saccharomyces cerevisiae* yeast strains used in this study were derivatives of BF264-15Daub (*ade1 his2 leu2*-*3,112 trp1*-*1a ura3*Δ*ns*), described previously [17]. Yeast cultures were grown at the indicated temperatures in YEP (1% yeast extract, 2% bacto-peptone, 0.005% adenine, 0.005% uracile) supplemented with 2% glucose (YEPD) or in selective minimal media. All strains were made isogenic by back crossing at least five times against our genetic background. The *siz1*::*KanMX4, siz2*::*KanMX4, cbf2*-*1*::*KanMX4* and *ctf13*-*30*::*KanMX4* strains were purchased at Euroscarf (Germany). The *bub2*Δ and *mad2*Δ mutants were kindly provided by Andrew Murray. The septin mutants were kindly provided by Michael McMurray. The *top2*-*SNM* strains as well as the *TOP2*-HA and *top2*-*SNM*-HA strains were kindly provided by Steve Elledge. The *rad17*Δ mutant was kindly provided by Errol Friedberg and the *mec3*Δ mutant by Maria-Pia Longhese. The *CDC13*-Myc_13_ strain was kindly provided by David Lydall. Constructs were made by using Polymerase Chain Reaction (PCR) to adapt the relevant restriction sites to the sequence of the gene and details of the constructs can be available upon request.

The viability of cells previously grown in liquid was determined by performing and analyzing the so-called “drop tests” or “spot assays”. To do this, cells from exponential growth cultures were counted with a hematocytometer and the cultures were then serially diluted by 1/10th and spotted onto YEPD (or selective medium) plates and incubated at the desired temperatures for 2-3 days before being photographed. In some cases, cells were just re-streaked onto YEPD plates and growth evaluated by visualizing the numbers and sizes of the growing colonies.

### 2.2. Construction of the ten1-sz, stn1-sz and cdc13-sz mutants

*stn1, ten1* and *cdc13* mutants were generated by PCR mutagenesis as described previously [17, 18, 46]. Briefly, *TEN1, STN1* and *CDC13* ORF flanked by sequences upstream of the ATG and post-STOP sequences were amplified by PCR (Dream Taq DNA polymerase, Fermentas) under mutagenic conditions in four distinct reactions, each reaction containing normal concentration of one of the four dNTPs, 0.2 mM, together with imbalanced and increased concentrations of the other three dNTPs (0.5 or 1.0 mM), as well as 3.0 mM MgCl2 (instead of 1.5 mM) and 0.5 mM MnCl2, in standard PCR buffer. Following a 30-cycle amplification, the PCR products were cleaned and, according to the gap repair method, were transformed into a double mutant strain harbouring both *siz1*Δ (::*KanMX4*) and either *stn1*::*TRP1, ten1*::*LEU2* or *cdc13*::*TRP1*, each surviving, respectively, owing to a *STN1*-*URA3, TEN1*-*URA3* or *CDC13*-*URA3* construct, together with a *STN1*-CEN-*LEU2, TEN1*-CEN-*TRP1* or *CDC13*-CEN-*LEU2* plasmid (CEN for centromeric), respectively, carrying flanking regions at each extremity previously cut at unique sites at (or close to) the ATG and stop codons to linearize the plasmids and allow recombination with the mutagenized PCR products. After *LEU2*^*+*^ or *TRP1*^+^ transformants plated out onto leucine- or tryptophan-lacking medium had developed at 24°C, they were replica-plated twice onto 5-FOA-containing medium at 24°C in order to force the loss of the *STN1*-*URA3, TEN1*-*URA3* or *CDC13*-*URA3* plasmid. Growing colonies were then pooled together and plated out onto YEPD plates at 24°C at various dilutions and, after colonies had developed, were replica-plated at 34 or 36°C. After comparing with the 24°C master plate, colonies that failed to grow at restrictive temperature were picked out and further expanded in liquid culture to perform plasmid recovery (from yeast to bacteria). Recovered plasmids were transformed back into the original mutant strains in order to confirm their temperature sensitivity. Further details concerning the mutagenic PCR processes are available upon request.

### 2.3. FACS analyses

For analysis of DNA content by flow cytometry, cells were fixed overnight, at 4°C, in 70% ethanol, treated with RNase (1 mg/ml) and pepsin (5 mg/ml), stained with propidium iodide (50 μg/ml) and analyzed in an Attune NxT Life Technologies acoustic focusing cytometer.

## 3. Results

### 3.1. Isolation of SIZ1 and TOP2 as extragenic suppressors of novel mutants of the telomeric Cdc13-Stn1-Ten1 complex

All three *Saccharomyces cerevisiae* CST genes, *CDC13, STN1* and *TEN1*, are essential genes. In the present study we set out to generate novel mutants of the CST complex with the objective of uncovering novel functions for this complex. Because numerous genetic screens have already been performed using mutants of *S. cerevisiae* CST, which may somewhat tend to some saturation, we used a different and unconventional strategy. A *cdc13*Δ *stn1*Δ *ten1*Δ triple mutant expressing the wild-type *CDC13*-*TEN1*-*STN1* fusion gene from a centromeric plasmid (see **Supplementary material**, sections 1.1 and 2.1 and **Fig. S1 A**) was perfectly viable (highlighting functionality of the *CDC13*-*TEN1*-*STN1* synthetic gene, referred to as *CST1* gene.

The *cdc13*Δ *stn1*Δ *ten1*Δ *p*-*CDC13*-*TEN1*-*STN1* strain was then used to construct temperature-sensitive mutants (see **Supplementary material** and **Fig. S1 A**). These so-called *cdc13*Δ *stn1*Δ *ten1*Δ *cst1* mutants (referred to as *cst*Δ *p*-*cst1*, to indicate the chromosomal mutation, followed by the covering plasmid), namely *cst*Δ *p*-*cst1*-*1, cst*Δ *p*-*cst1*-*4, cst*Δ *p*-*cst1*-*5* and *cst*Δ *p*-*cst1*-*15* (**Fig. S1 B**) were then used in genetic screens to isolate suppressors of their temperature sensitivity (see **Supplementary material**, sections 1.2 and 2.2 and **Fig. S2**). *SIZ1*, which codes for a SUMO E3 ligase [47, 48] and *TOP2*, which codes for topoisomerase II (which, interestingly, is sumoylated by Siz1, as well as by Siz2, a related SUMO E3 ligase [49] were isolated as extragenic suppressors of these *cst*Δ *p*-*cst1* mutants (**Fig. S2**). The relevance of these genetic interactions was suggested by the existence of negative genetic interactions between the *stn1*-*13* and the *siz1*Δ mutants in a genome-wide screen [50]. In this type of experiments, a rescue phenotype upon overexpression of *SIZ1* or *TOP2* means improved growth capacity and increased survival in face of this telomeric damage.

Interestingly, in the same screens on the *cst*Δ *p*-*cst1* mutants, *TEN1* was frequently found to rescue all four *cst*Δ *p*-*cst1* mutants (a total of 12 genomic fragments isolated; **Fig. S2 B**), but *CDC13* and *STN1* were never isolated. Overexpression of *CDC13* or *STN1* alone from a similar multicopy plasmid to those from the genomic libraries did not rescue the *cst*Δ *p*-*cst1* mutants (**data not shown**), thus suggesting that the failure to isolate either one of these genes in the genetic screens was not due to the fact that those two genes were simply missing from the libraries. Furthermore, genomic fragments from both libraries containing either *STN1* or *CDC13* have been isolated by us in the past in separate projects. The rescue of *cst*Δ *p*-*cst1* by *TEN1* found here suggests that the particular conformation adopted by the *CDC13*-*TEN1*-*STN1* fusion polypeptide mutants prevents Ten1 (perhaps “blocked” between Cdc13 and Stn1) from accomplishing an essential function that can be provided by YEp-*TEN1* expression.

### 3.2. Isolation of ten1-sz, stn1-sz and cdc13-sz mutants on the basis of rescue by overexpression of SIZ1

In addition to the *cst*Δ *p*-*cst1* mutants rescued by *SIZ1* overexpression, it was important to also isolate single mutants for each of the three CST subunits to confirm that the effects observed were not specific to the triple fusion protein, and to simplify further studies by the use of more classical single gene mutants. Of note, many previously described mutants of CST, such as *ten1*-*31, ten1*-*16, stn1*-*13, stn1*-*101, stn1*-*154* and *cdc13*-*1* (see references [15, 17, 18, 51]) were not rescued by *SIZ1* overexpression (**data not shown**). Therefore, it was most important to design a strategy to isolate individual mutants for each of the three CST genes in order to elucidate these potentially new CST functions.

We thus generated, by PCR mutagenesis (see **Materials and methods** ; section 2.2), temperature-sensitive mutants of *CDC13, STN1* and *TEN1* in a *SIZ1*-deleted background, reasoning that this might increase the chances to generate mutants more specifically defective in Siz1-related functions - the idea was that a *ten1 siz1*Δ, a *stn1 siz1*Δ or a *cdc13 siz1*Δ strain would be more prone to generate temperature-sensitive clones than the corresponding *SIZ1*^+^ strains, and that these would be more specifically defective in Siz1-related functions (**Fig. 1 A**, top panels). This is exactly what happened. Indeed, in the case of *stn1 siz1*Δ, only three temperature-sensitive clones (named *stn1*-*sz* mutants, *sz* referring to Siz1 implication in *stn1* phenotypes) were isolated from approximately 20,000 transformants. The *stn1*-*sz1* no longer rescued the original strain upon re-transformation, but *stn1*-*sz2* and *stn1*-*sz3* passed this test. Interestingly, the temperature sensitivity of both, higher for *stn1*-*sz2* than for *stn1*-*sz3*, could be rescued by overexpressing *SIZ1*, clearly visible on the increased capacity to form colonies, as well as on the partial suppression of morphological defects (**Fig. 1 B** for *stn1*-*sz2, stn1*-*sz3* not shown). Using the same approach, two mutants of *CDC13* (*cdc13*-*sz2* and *cdc13*-*sz23*), selected among thirty-three temperature-sensitive clones isolated from around 5,000 transformants, could be rescued by *SIZ1* overexpression. Similarly, the temperature-sensitivities of three *TEN1* mutants (*ten1*-*sz5, ten1*-*sz8* and *ten1*-*sz12*) selected among thirteen temperature-sensitive clones in a total of around 10,000 transformants), were partially suppressed at 34° and 36°C upon overexpression of *SIZ1* (**Fig. 1 B** and **data not shown**). Sequences of all *cst*-*sz* mutants are provided in **Figure 1C**.

**Figure 1.**
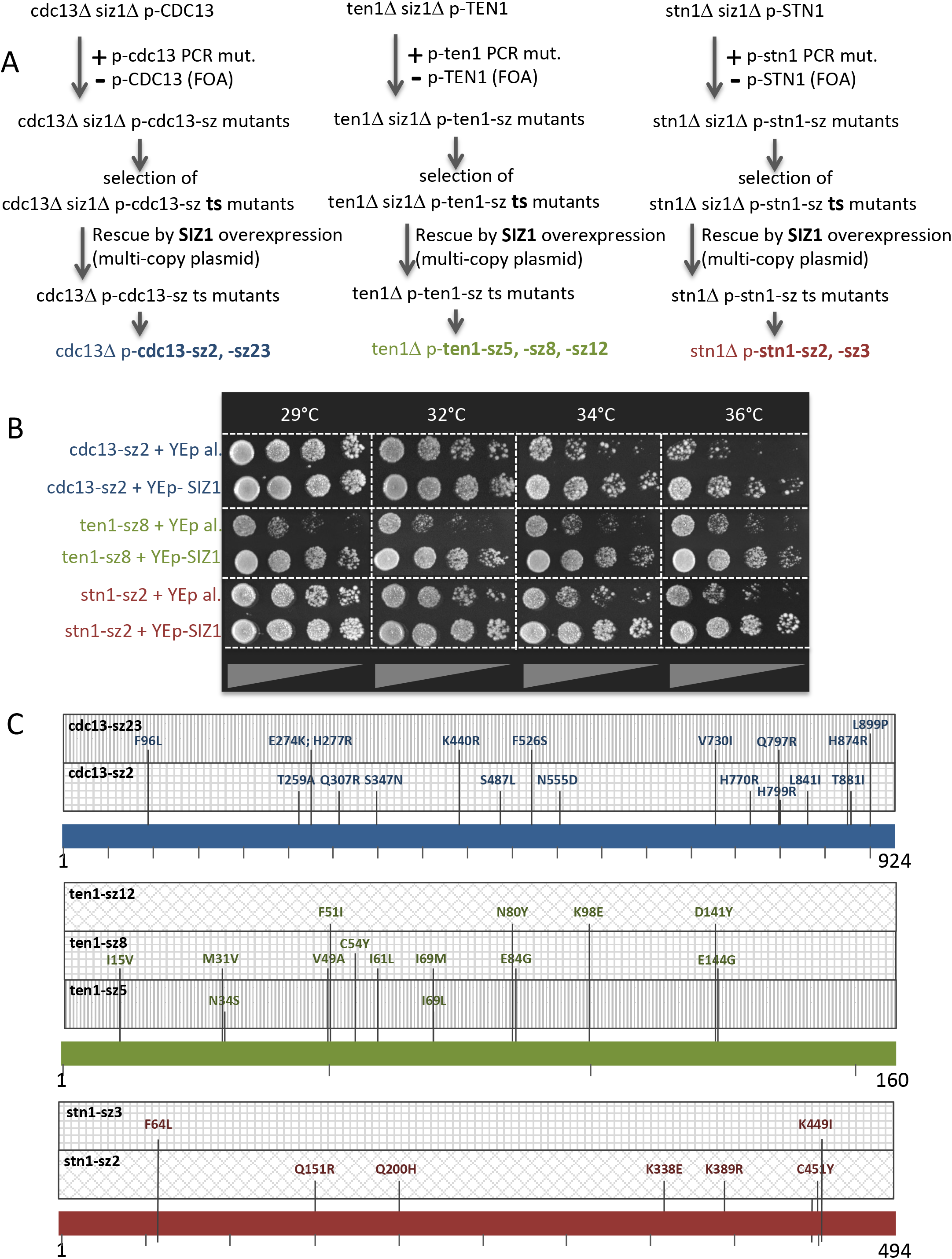
Isolation and characterization of *cdc13*-*sz, stn1*-*sz* and *ten1*-*sz* mutants sensitive to intracellular Siz1 dosage. (**A**) Schematic protocol used to generate temperature-sensitive *cdc13, ten1* and *stn1* mutants in a *siz1* null background (*siz1*Δ) by PCR mutagenesis coupled to gap repair (see Materials and methods, section 2.2), followed by selection of mutants that can be rescued by overexpression of *SIZ1* from a 2μ multi-copy plasmid. Two *cdc13*Δ *p*-*cdc13*-*sz* mutants (hereafter referred to as *cdc13*-*sz2* and *cdc13*-*sz23*), three *ten1*Δ *p*-*ten1*-*sz* mutants (*ten1*-*sz5, ten1*-*sz8* and *ten1*-*sz12*) and two *stn1*Δ *p*-*stn1*-*sz* mutants (*stn1*-*sz2* and *stn1*-*sz3*) were isolated. (**B**) Rescue of the temperature-sensitive growth defects of the *cdc13*-*sz2, ten1*-*sz8* and *stn1*-*sz2* mutants upon overexpression of *SIZ1* from a 2μ multi-copy vector (+ YEp-SIZ1), with expression of plasmid alone (+ YEp al.) as a control. (**C**) Sequences of all Cdc13-sz, Ten1-sz and Stn1-sz mutant proteins.

All *cst*-*sz* mutants tested, namely *stn1*-*sz2 siz1*Δ, *cdc13*-*sz2 siz1*Δ, *cdc13*-*sz23 siz*1Δ and *ten1*-*sz8 siz1*Δ, grew no worse than their *SIZ1*^+^ counterparts, except perhaps the *ten1*-*sz8 siz1*Δ mutant (**data not shown**). Nevertheless it is clear that overexpression of *SIZ1* rescued these mutants, pointing to an involvement of Siz1 in their loss of function. Therefore, these *cdc13*-*sz, stn1*-*sz* and *ten1*-*sz* mutants appear to be sensitive to intracellular Siz1 dosage.

### 3.3. Role of the three major G2/M checkpoints in the detection of damage in the stn1-sz2 mutant

Our next objective was to determine whether or not the damage in the *cst1*-*sz* mutants was activating one or more of the three major G2/M checkpoints, namely the Mec1-Mec3 DNA damage checkpoint [52], the spindle orientation checkpoint [53] and the spindle assembly checkpoint [54]. All three major G2/M checkpoints played a role in the detection of *stn1*-*sz2* defects as attested by visible effects on viability when the DNA damage checkpoint (*rad17*Δ) or of one of the two spindle checkpoints (*mad2*Δ and *bub2*Δ) had been genetically inactivated (**Fig. 2 A**). Interestingly, two different effects of these checkpoint mutations on *stn1*-*sz2* growth were observed. Thus, inactivation of *RAD17* or of *MAD2* in the *stn1*-*sz2* mutant at 34-36°C resulted in an increase in cell viability, indicated by improvement of colony formation or colony size (**Fig. 2 A**). This phenotype was reminiscent of that described for the *cdc13*-*1 rad17*Δ mutant and is, in fact, characteristic of forcing passage of the checkpoint-mediated cell cycle arrest [55, 56]. This same phenotype has also been described for the *cdc13*-*1 bub2*Δ mutant [57] and interpreted the same way as that of the *cdc13*-*1 rad17*Δ mutant [55, 56]. On the other hand, the phenotype of the *stn1*-*sz2 bub2*Δ double mutant was more classical, exhibiting a strong synthetic lethality indicative of the dramatic consequences of preventing activation of the Bub2 checkpoint in the presence of *stn1*-*sz2* damage (**Fig. 2 A**).

**Figure 2.**
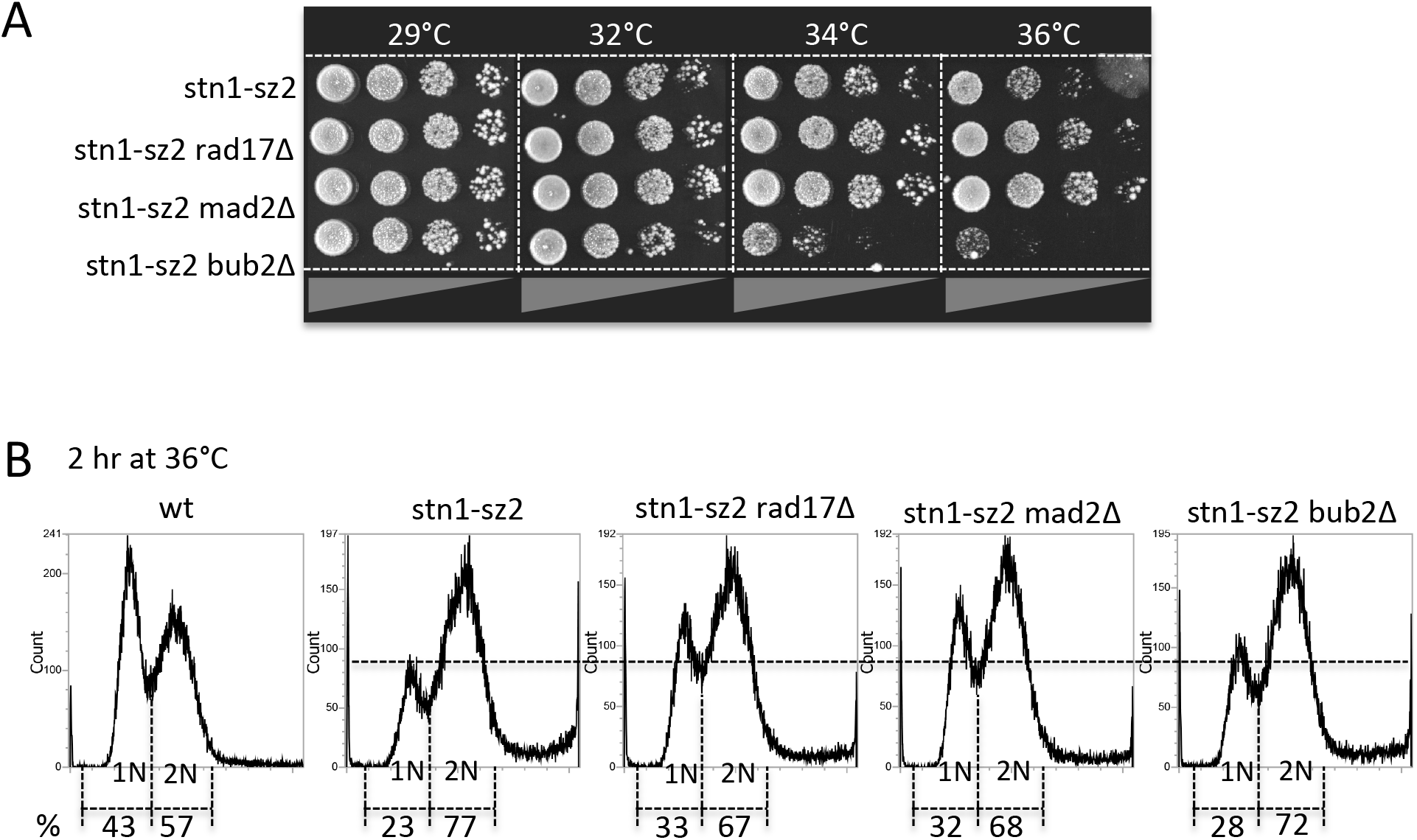
Effects on the temperature-sensitive *stn1*-*sz2* mutant of genetic inactivation of the G2/M DNA damage checkpoint (*rad17*Δ mutation) or of one the two spindle checkpoints (*mad2*Δ and *bub2*Δ mutations) on cell viability (**A**) and on cell cycle distribution assessed by FACS analysis (**B**).

Analysis of DNA content by flow cytometry (FACS) confirmed the implication of the Mec1-dependent DNA damage and Mad2-dependent spindle checkpoints in the arrest of the stn1-sz2 mutant at the restrictive temperature of 36°C (**Fig. 2 B**). On the other hand, the *stn1*-*sz2 bub2*Δ mutant cells were distributed along the different cell cycle stages just like the *stn1*-*sz2* cells, thus confirming that synthetic lethality rather than deregulation of cell cycle arrest was taking place. In summary, the *stn1*-*sz2* mutant at 36°C produces damage that normally triggers cell cycle arrest at G2/M and we observe that complete genetic inactivation of either *RAD17* or *MAD2* prevents that arrest, while inactivation of *BUB2* in *stn1*-*sz2* kills the cells. Thus, all three G2/M checkpoint pathways detected damage produced in the *stn1*-*sz2* mutant.

### 3.4. Looking for the sumoylated targets of Siz1 in stn1-sz2

As seen above, sumoylation appears to play some role in the rescue of the CST mutants described in this study. At least two likely candidates emerged as being possibly involved in this, the septins and Top2.

We found that Top2 was sumoylated in the *stn1*-*sz2* mutant (**Supplementary Results**, section 2.3 and **Fig. S3 A**), and that Cdc3 septin was also sumoylated in this mutant, as well as in the *ten1*-*sz* and *cdc13*-*sz* mutants (**Fig. S3 B, C** and **Fig. S4**). However, in neither case could we establish whether sumoylation of Top2 and Cdc3 resulted from the damage inflicted by the mutations in CST, or were rather due to the fact that mutant cells were arrested in mitosis as a consequence of a cell cycle effect of the temperature-sensitive mutations.

Previous studies have established septins as major targets of Siz1 SUMO ligase activity [47, 48, 58, 59]. Among the five *S. cerevisiae* septins functioning during vegetative growth, Cdc3 is the principal target of Siz1-dependent sumoylation (four lysine residues identified), followed by Shs1 (two lysine residues) and Cdc11 (one lysine residue) [48, 58]. In a SUMO interactome study, all five septins (Cdc3, Cdc10, Cdc11, Cdc12 and Shs1) were found to bind Siz1 [60]. In addition, in a proteome-wide study, Cdc12, as well as Cdc3, Cdc11 and Shs1, were also found to be sumoylated [61]. Interestingly, negative genetic interactions between *TEN1* and *CDC12* were uncovered in a previous study [62]. Here, we found that cell viability (**Fig. 3 A**) and morphological (**data not shown**) defects were much aggravated when the *ten1*-*31* mutation and a mutation in a septin gene were combined, compared with the single mutants alone (**Fig. 3 A**). This contrasts with two well-documented mutant alleles of *CDC13* and *STN1, cdc13*-*1* [15] and *stn1*-*13* [17], which exhibit moderate genetic interactions with a *cdc10* (but not *cdc11* or *cdc12*) and a *cdc12* (but not *cdc10* or *cdc11*) mutant, respectively (**not shown**). Importantly, stronger negative genetic interactions were observed between the septin mutants and the *stn1*-*sz2* mutant (**Fig. 3 B**) than between the septin mutants and the classical *stn1*-*13* and *cdc13*-*1* mutants. Thus these new *cst1*-*sz* mutants have a loss of function that is more sensitive to the presence of intact septins than the mutants of CST already described in the literature.

**Figure 3.**
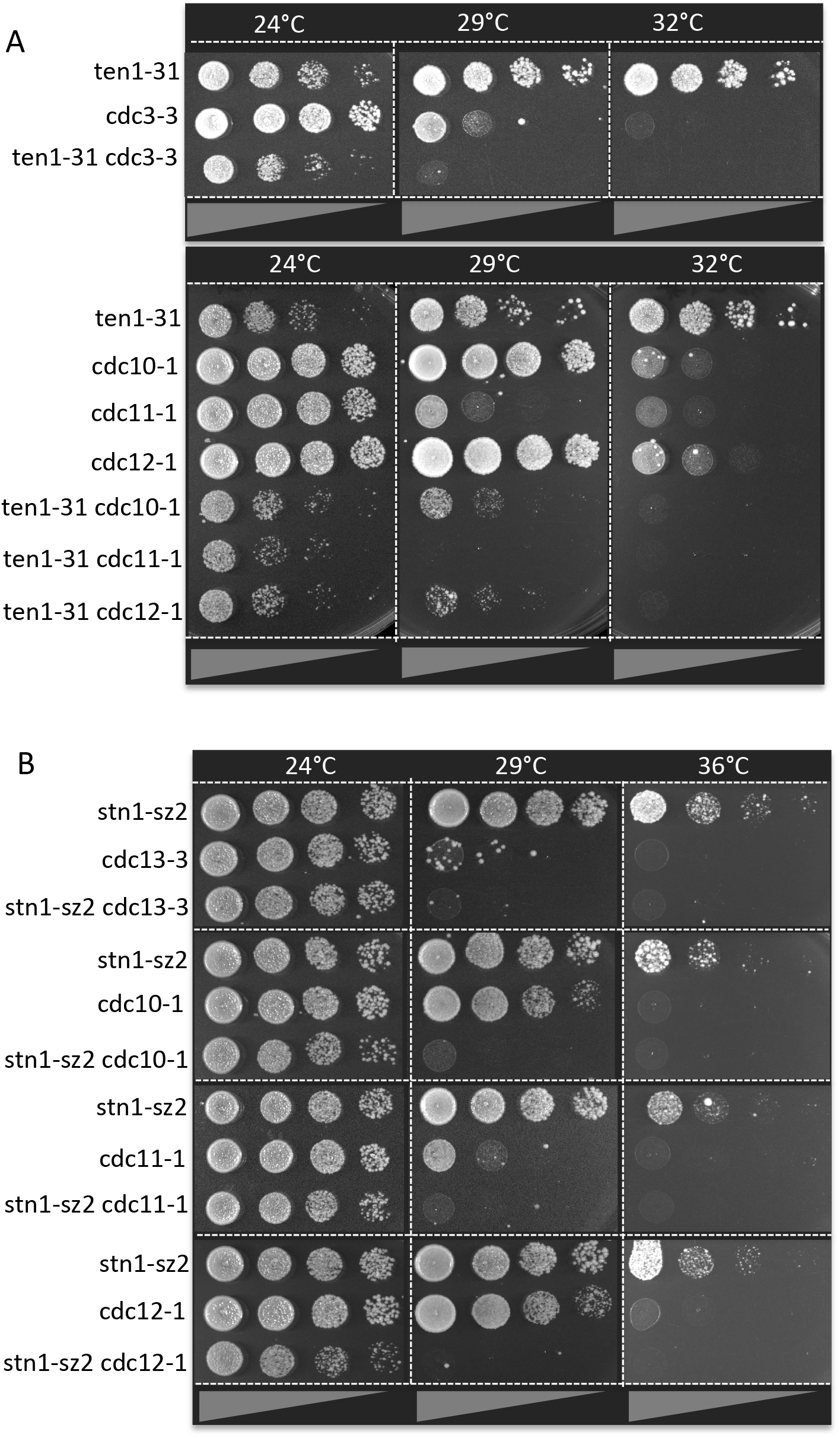
Genetic interactions between the Cdc13-Stn1-Ten1 complex and septins. **(A)** The temperature-sensitive *ten1*-*31* mutant exhibited negative genetic interactions (also frequently referred to as synthetic growth defects) with the indicated temperature-sensitive septin mutants. (**B**) The temperature-sensitive *stn1*-*sz2* mutant exhibited strong genetic interactions with several temperature-sensitive mutants of septin, as indicated.

The sumoylated form of Top2 represents a metaphase-anaphase checkpoint both in yeast and humans [63] and Siz1 is necessary for most of Top2 sumoylation [49]. To check whether sumoylation of Top2 was involved in the rescue of CST loss of function by *SIZ1*, we used a mutant of *TOP2, top2*-*SNM* (Sumo No More). *top2*-*SNM* encodes a protein with six mutations in lysine residues sumoylated by SUMO/Smt3 [64]. Experiments with the *top2*-*SNM* mutant suggested that Top2 sumoylation might play some role in these mechanisms of rescue of *stn1*-*sz2* by *SIZ1* overexpression (**Fig. 4 A**) and that overexpression of *top2*-*SNM* was less efficient than that of *TOP2* in rescuing the *cst*Δ *p*-*cst1*-*1* mutant (**Fig. 4 B**).

**Figure 4.**
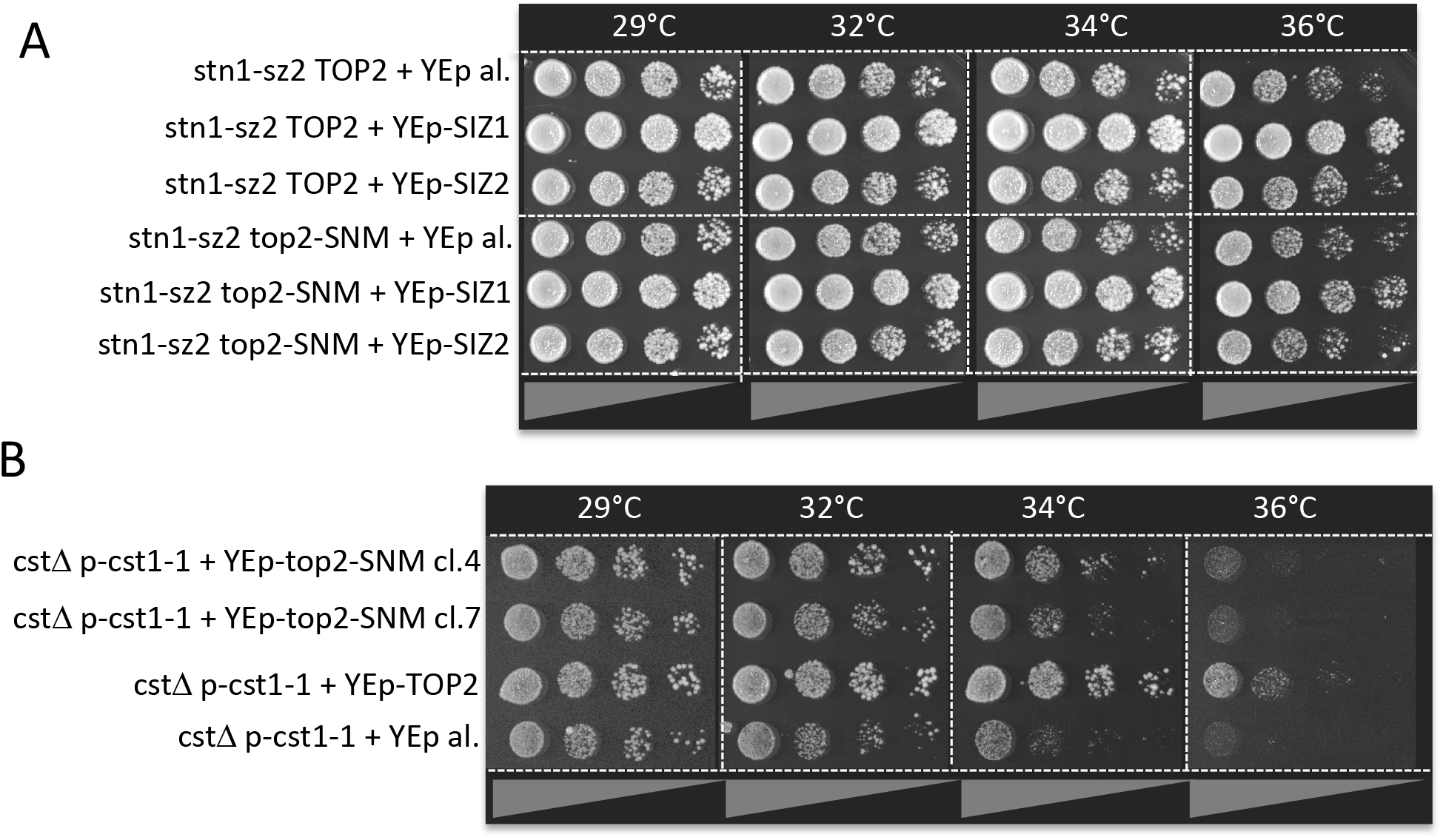
Effects of Top2 sumoylation on the rescue of the *stn1*-*sz2* and *cst*Δ *p*-*cst1*-*1* mutants.(**A**) Rescue of the *stn1*-*sz2* mutant by overexpression of *SIZ1* (or of *SIZ2* or of vector alone; all from a 2μ multi-copy plasmid, “YEp-SIZ1”, “YEp-SIZ2 or “YEp al.”) was less efficient when *top2*-*SNM* mutations were expressed from *TOP2* locus, rendering Top2 unsumoylatable, but was nevertheless still visible. (**B)** Overexpression of *top2*-*SNM* (from a 2μ multi-copy plasmid) was a little less efficient than that of *TOP2* in rescuing *cdc13*Δ *stn1*Δ *ten1*Δ *p*-*cst1*-*1* (two clones of *top2*-*SNM* constructs, “cl.4” and “cl.7”, are shown here).

## 4. Discussion

In the present study we have isolated novel mutants of the conserved telomeric Cdc13-Stn1-Ten1 complex (CST) of the yeast *Saccharomyces cerevisiae*. These mutants are unique in the way the damage they generate simultaneously activates both major G2/M spindle checkpoints together with the G2/M DNA damage checkpoint. In cancer research a big challenge is to accurately identify the origin and nature of the damage generated at the telomeres and the way this is recognized by the checkpoint machineries, which in turn will affect cell cycle progression in the tumor cells. Thus, understanding the mechanisms by which telomeric damage controls the cell cycle is of major importance in cancer biology, as well as in aging-related (and other) telomeropathies [6, 8, 65]. The conserved CST (CTC1-STN1-TEN1) telomeric complex plays major roles in these pathways, particularly by providing telomere end protection and regulation of telomerase accessibility [10, 11, 27, 31, 66].

### 4.1. Isolation of mutants of CST rescued by overexpression of SIZ1 or TOP2

The new mutants of CST isolated here represent invaluable tools to study (a) yet uncharacterized function(s) of CST. Such mutants with a loss of function partially rescued by *SIZ1* or *TOP2* have not been previously uncovered, the unconventional strategy of genetic screening designed here to isolate these CST mutants presumably introducing a bias towards isolation of this particular class of CST mutants. A particular, unnatural, conformation adopted by the mutagenized *cdc13*-*ten1*-*stn1* hybrid gene (that we call the *CST1* gene) might have favoured the accumulation of unusual mutations.

The main conclusion of the present study is that defects in *S. cerevisiae* CST are simultaneously sensed not only by the G2/M DNA damage checkpoint, but also by the two major G2/M spindle checkpoints. Most of the mutants of CST identified to date activate the Mec1-mediated DNA damage checkpoint [52], due to the fact that most of them are compromised in telomeric DNA binding and accumulate abnormal levels of single-stranded telomeric DNA, in budding yeast [15, 46] and other organisms [11, 31, 67]. The Bub2-dependent checkpoint monitors correct orientation of the mitotic spindle [53], while the Mad2-dependent checkpoint monitors attachment of the kinetochores (centromeric structures of the chromosomes) to the spindle microtubules [54]. It should be noted that the *cdc13*-*1* mutant has previously been reported to activate the Bub2 spindle orientation checkpoint [57]. While the pathways at the origin of this genetic interaction have not yet been elucidated, it is tempting to speculate that they might correspond to those suspected to be deregulated in our new mutants of CST.

In addition to *SIZ1* and *TOP2, TEN1* (but not *STN1* and *CDC13*) was also a good suppressor of the *cst*Δ *p*-*cst1* mutants, perhaps highlighting the fact that essential domains of Ten1 are masked by the surrounding Cdc13 and Stn1 in the synthetic fusion protein. In this respect, it would be interesting to construct temperature-sensitive hybrid fusion CST genes with other orders of arrangement of the three genes within the fusion, to see whether their subsequent mutagenesis also leads to isolation of *SIZ1* and *TOP2* extragenic suppressors. In addition, structural modelling of the fusion encoded by the synthetic *CST1* gene might allow to pinpoint particular losses of interactions with other mitotic and/or telomeric actors when compared with structural data from each of the single CST subunits.

Interestingly, mutagenesis of each of the single CST genes, *CDC13, STN1* and *TEN1*, under conditions of complete depletion of intracellular Siz1 (in a *siz1*Δ background), also introduced a bias in the isolation of temperature-sensitive mutants that could be rescued by overexpression of *SIZ1*. Indeed, under such conditions (*siz1*Δ background), it became much easier to isolate temperature-sensitive alleles than under normal conditions, as shown in our genetic screenings. These so-called *cdc13*-*sz, stn1*-*sz* and *ten1*-*sz* might be more amenable to future structural and genetic studies, as the characteristics of the mutant proteins might be directly compared to those of the corresponding wild type proteins.

Another finding of the present study is to have generated a novel, artificial, fusion hybrid gene that is fully functional in terms of complementing the *cdc13*Δ *stn1*Δ *ten1*Δ triple deletion mutant. We refer to this synthetic gene as the *CST1* gene. We chose to name this synthetic gene *CST1* rather than *CTS1* (which would have been more logical given the order of the genes in the fusion) because *CTS1* is an already deposited gene name (encoding endochitinase) and, in addition, because CST is the name of the complex composed of Cdc13, Stn1 and Ten1 and the name cst1 for these mutants is therefore more descriptive of their functions. This functional synthetic *CST1* gene will be a useful tool in future studies aiming to further decipher the structural interactions of the three subunits with other telomeric proteins.

### 4.2. Mechanisms of rescue of mutants of CST by overexpression of SIZ1 or TOP2

The isolation of *SIZ1* and *TOP2* as extragenic multicopy suppressors of the newly described *cst1* mutants led us to examine the possibility that SUMO post-translational modifications [44, 68-70] might be involved in these mechanisms of rescue. We found here that rescue of CST mutants by overexpression of *TOP2* somewhat depended on Top2 sumoylation, in agreement with the sumoylated form of Top2 representing a metaphase-anaphase checkpoint both in yeast and humans [63]. Several additional arguments suggested that sumoylation of the septins was also possibly involved in the mechanisms of rescue of the new cst1 mutants by overexpression of *SIZ1*, but this remains to be demonstrated.

Given the diversity of mechanisms of extragenic rescue of mutations described in the literature (see **Supplementary Discussion**, section 3.1), future studies will be needed for a complete understanding of how exactly overexpression of *SIZ1* or of *TOP2* leads to diminished damage and increased viability in these new mutants of CST.

Since the new mutants of *S. cerevisiae* CST described here activate the two major spindle checkpoints, physical association of hCST with cohesin [29] might be pertinent to the mechanisms studied here in yeast. In addition, we need to understand how septin sumoylation can affect cell cycle progression in response to damage, such mechanisms becoming pertinent to study following the recent finding that a defect in septin sumoylation in humans correlates with defects in cytokinesis [71].

### 4.3. Conclusions

The main conclusion of the present study is that defects in *S. cerevisiae* CST (Cdc13-Stn1-Ten1) are sensed not only by the G2/M DNA damage checkpoint, but also by the two major G2/M spindle checkpoints. The pathologies associated with mutations in human CTC1 and STN1, mainly dyskeratosis congenita and Coats Plus ([72] and references therein), might therefore stem not only from deregulation in CTC1-STN1-Pol-α-telomerase interactions [73], but also from potential defects in mitotic spindle stability conferred by the mutations in *S. cerevisiae* CST described here. The present data suggest that septins might represent a potential target of the spindle checkpoints that become activated after telomeric damage in these new mutants of CST has been generated.

## Supporting information

Supplematay materia text and figures

## Supplementary Materials

Supplementary data associated with this article can be found in the online version at www.mdpi.com/xxx/s1,

## Author Contribution

Grandin Nathalie and Charbonneau Michel: conceived the experiments, acquired funding, conducted the experiments, analyzed the results, wrote the manuscript.

## Acknowledgments

The authors thank the LIGUE Nationale contre le Cancer (CCAURA, Comités Puy-de-Dôme et Allier) for its financial support. We thank Michael McMurray, Andrew Murray, Steve Elledge, Errol Friedberg, Maria-Pia Longhese and David Lydall for the gifts of strains and plasmids. We also thank Charles White for helpful discussions and critical reading of the manuscript.

