## Supplementary material for "Dysfunction of telomeric Cdc13-Stn1-Ten1 simultaneously activates the DNA damage and spindle checkpoints": Supplematay materia text and figures

#### 1. Materials and methods

*1.1. Construction of temperature-sensitive mutants of a synthetic* CDC13-TEN1-STN1 *hybrid gene*

First, a low-copy plasmid expressing the *CDC13-TEN1-STN1* in-frame fusion gene was constructed by cloning, in a centromeric plasmid (YCplac33, *URA3* marker; YCplac series; 2 to 4 copies, [1]), the entire ORF of the first part of the hybrid gene, *CDC13*, plus upstream promoter sequences of 300 nt, in front of *TEN1* ORF (and in reading-frame with it), itself in front of *STN1* ORF (and in reading-frame with it), which included its stop codon, followed by 120 nt of *STN1*'s post-stop natural sequences. Next, a *cdc13*Δ (*cdc13*::*URA3* reverted to ura- on 5-FOA, a drug used to counterselect *URA*+ cells) *stn1*Δ (*stn1*::*TRP1*) *ten1*Δ (*ten1*::*KanMX4*) triple mutant diploid strain was transformed with the wild-type *CDC13-TEN1-STN1* fusion gene (*URA3* marker) described above and sporulated to allow isolation of a *cdc13*Δ *stn1*Δ *ten1*Δ *CDC13-TEN1-STN1* haploid strain. This strain was perfectly viable (highlighting functionality of the *CDC13-TEN1-STN1* synthetic gene, referred to as *CST1* gene) and did not exhibit any apparent morphological defects at temperatures comprised between 24 and 36°C (**Fig. S1**). Next, the *cdc13*Δ *stn1*Δ *ten1*Δ YCplac33-*CDC13-TEN1-STN1* triple mutant strain (*URA3* marker) was transformed with mutagenized YCplac111-*cdc13-ten1-stn1* plasmid (*LEU2* marker). To do this, the YCplac111-*CDC13-TEN1-STN1* plasmid was cut with *Stu*I and genecleaned to remove most of the *CDC13-TEN1-STN1* insert (see below), together with PCR-mutagenized *cdc13-ten1-stn1* fusion reactions that encompassed the three genes (*CDC13-TEN1-STN1*) from 300 nt upstream of *CDC13*'s ATG to 134 nt downstream of *STN1*'s stop codon, a process known as gap repair-coupled PCR mutagenesis. *Stu*I cut only twice in the YCplac111-*CDC13-TEN1-STN1-LEU2* plasmid, once at 105 nt upstream of *CDC13*'s ATG and a second time at 44 nt downstream of *STN1*'s stop codon. This left 196 nt of *CDC13*'s promoter sequences and 90 nt of *STN1*'s post-stop sequences for homologies with the mutagenic PCR reactions to allow the repair of the StuI-gapped YCplac111-*CDC13-TEN1-STN1-LEU2* plasmid using the co-transformed *cdc13-ten1-stn1* mutagenic PCR reactions. Dream Taq DNA polymerase (Fermentas) was used for mutagenic PCR under conditions of imbalanced and increased concentrations of the dNTPs, together with the presence of 3.0 mM $MgCl_2$ (instead of 1.5 mM) and 0.5 mM $MnCl_2$, in standard PCR buffer. Following selection of the transformants on -Leu selective medium, the *CDC13-TEN1-STN1* fusion wild-type gene was forced to extrusion following growth on 5-FOA to counterselect *URA*+ cells (**Fig. S1**; further details concerning the mutagenic PCR processes are available upon request). After several successive experiments in which the 24°C master plates were compared with replicas incubated at 36°C, we could isolate four temperature-sensitive mutants, namely *cst*Δ (for *cdc13*Δ *stn1*Δ *ten1*Δ) *p-cst1-1*, *cst*Δ *p-cst1-4*, *cst*Δ *p-cst1-5* and *cst*Δ *p-cst1-15* (**Fig. S1**). These mutants were further expanded in liquid culture to perform plasmid recovery (from yeast to bacteria) and recovered plasmids were transformed back into the original mutant strains in order to confirm their temperature sensitivity and avoid any possible contribution of other mutations in the strains.

*1.2. Isolation of extragenic suppressors of the* cdc13Δ stn1Δ ten1Δ p-cst1 *mutants*

To isolate extragenic suppressors of the temperature-sensitive *cst*Δ *p-cst1* mutants described above, cells were transformed with genomic DNA libraries expressing high copy-plasmids, namely the pFL-44L [2] and YEp24 libraries [3] both 2μ and *URA3*-marked. Clones that grew better than original mutants at 32 or 34°C were selected following replica-plating on -Ura drop-out plates by comparing with the master plate incubated at 24°C. Plasmids were then recovered from the selected clones (plasmid recovery from yeast to bacteria), distinguishing every time, by restriction enzyme analysis, the rescuing library plasmid from the mutant *cdc13-ten1-stn1* plasmid (because both plasmids were present in the same cells). Bank plasmids were then transformed back into the original mutants to re-assess rescue of the temperature sensitivity. Selected bank plasmids were then sequenced and restricted by separately cloning all the genes contained in the genomic DNA fragment (all of a size comprised between around 4.0 and 8.0 Kb) and all of these genes were transformed separately into the original corresponding *cst*Δ *p-cst1* mutant to assess its capacity to rescue the temperature-sensitive defect.

*SIZ1* (YDR409w) ORF (2,715 nucleotides located between positions 1,289,406 and 1,292,120 of chromosome IV), contained in a YEp24 genomic bank plasmid that comprised a 4,084 nucleotides-long fragment of sequences located between nucleotides 1,288,536 and 1,292,620 of chromosome IV, rescued the *cst*Δ *p-cst1-1* mutant. *TOP2* (YNL088w) part (only the last 2,716 nucleotides of the 4,287 nucleotides-long ORF comprised between positions 459,274 and 461,990 of chromosome XIV), contained in a 4,464 nucleotides-long fragment of pFL-44L genomic bank plasmid comprising nucleotides 459,274 to 463,738 of chromosome XIV, rescued the *cst*Δ *p-cst1-15* mutant. We note that M524 of TOP2 ORF precisely coincides with the beginning of the genomic fragment that rescued the *cst1-15* mutant and, therefore, we surmise that the last 905 amino acids of Top2 (out of a total of 1429) were expressed under the control of M524 as the initiating methionine. In addition, *TEN1* contained in YEp24 and pFL-44L libraries was frequently found to rescue all four *cst*Δ *p-cst1* mutants (a total of 12 genomic fragments isolated), but *CDC13* and *STN1* were never isolated.

##### *1.3. Sumoylation of Top2 and septin*

Septin sumoylation was visualized following Western blotting (using anti-HA monoclonal antibody) of whole TCA (trichloroacetic acid) extracts prepared from asynchronous wild-type cells expressing $HA_2$-*CDC3* from *CDC3* genomic locus and run on polyacrylamide gels. To visualize the sumoylated forms of septins, it was necessary that the isopeptidases that cleave SUMO off its substrates virtually instantly when cells are lysed under native conditions, be completely inhibited, which was achieved by preparing TCA extracts [4]. To do this, cell pellets were resuspended in 20% TCA, mechanically broken with glass beads and after the latter were discarded by centrifuging at 1,000 rpm, the suspension was diluted in 5% TCA in order to obtain a 10% TCA suspension which was immediately centrifugated at 3,000 rpm. The resulting protein pellet was resuspended in loading buffer and boiled prior to separation by SDS-PAGE gel electrophoresis.

Sumoylation of the Cdc3 septin was measured in denaturing TCA extracts, as described above, as the sumoylated bands have been well described in the literature [4-6]. However, to measure Top2 sumoylation, identification of the sumoylated bands required immunoprecipitation. Usually, experiments aiming to detect sumoylation on a particular protein are performed using Ni-chromatography (with 6 His tag) under denaturing conditions, 6 M guanidium chloride. Conventional IP (immunoprecipitation) is usually not denaturing enough to allow total inhibition of SUMO isopeptidases (Ulp1 and Ulp2), which disrupt the isopeptide bond between Smt3 (SUMO) and the sumoylated protein in the cell extract. Yet, here, we performed IP of Top2-HA2 with monoclonal anti-HA and, in parallel in the same extracts, co-IP between Top2-HA2 and Myc2-Smt3, using also monoclonal anti-Myc antibody. Strains were tagged at the corresponding genomic loci to express simultaneously endogenous $HA_2$-*TOP2* and $Myc_2$-*SMT3*). This can work fine only when the concentration of detergents in the RIPA buffer is compatible with both efficient co-IP and preservation of SUMO binding to Top2.

In all experiments, results were analyzed with a BioRad ChemiDoc Imaging System (GEHealthcare) and quantified using the Image Lab software (BioRad).

##### *1.4. Telomere length measurements*

Telomere length measurement using the classical $^{32}$P-based telomere restriction fragment analysis method (TRF) was performed as previously described [7]. Briefly, genomic DNAs were prepared, separated in a 0.9% agarose gel (in TBE) run in TBE buffer overnight and, after denaturation, transferred and hybridized with a 270 base pair $TG_{1-3}$ $^{32}$P-labeled telomeric probe. Following digestion of genomic DNA with *Xho*I, to cut within the Y' regions of chromosomes, telomere tracts of wild-type cells appear as a broad band of ~ 1.1-1.2 Kb which contains the most terminal 300-350 nt of $TG_{1-3}$telomeric sequences and therefore represents the average length of the telomeres. Results were analyzed using an AmershamTM Typhoon IP phosphorimager (GE Healthcare) and the ImageQuant TL software.

#### 2. Results

##### *2.1. Construction of novel mutants of the telomeric Cdc13-Stn1-Ten1 complex*

All three *Saccharomyces cerevisiae* CST genes, *CDC13*, *STN1* and *TEN1*, are essential genes. In the present study we set out to generate novel mutants of the CST complex with the objective of uncovering novel functions for this complex. Because numerous genetic screens have already been performed using mutants of *S. cerevisiae* CST, which may somewhat tend to some saturation, we used a different and unconventional strategy. Under current conditions it is difficult to isolate extragenic suppressors of temperature-sensitive *cdc13* mutants other than *STN1* and, likewise, difficult to isolate extragenic suppressors of temperature-sensitive stn1 mutants other than *TEN1*. Empirically we reasoned that a genomic fragment of a given DNA library containing either one of the three CST

Figure S1

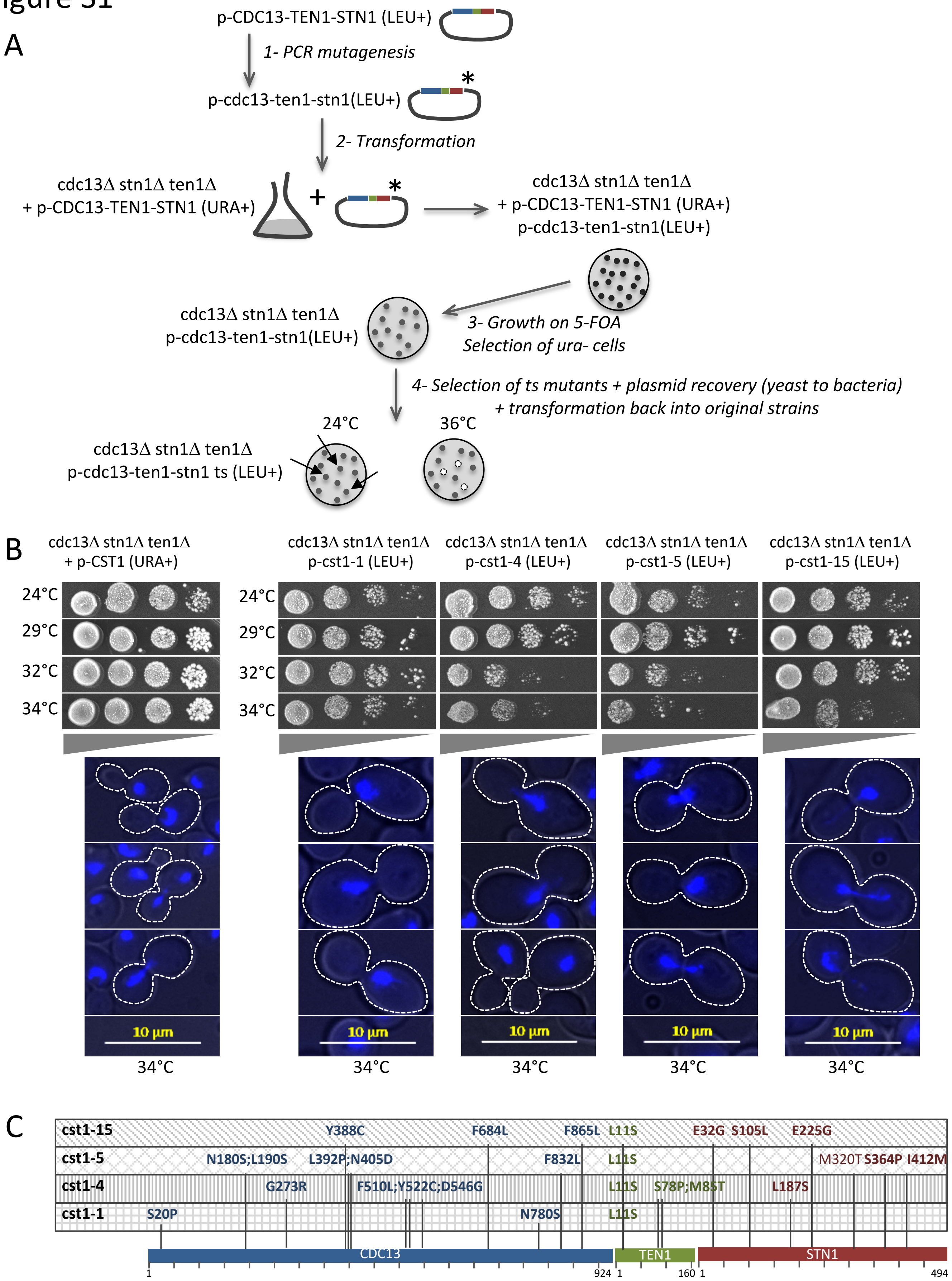
A
p-CDC13-TEN1-STN1 (LEU+)
1- PCR mutagenesis
p-cdc13-ten1-stn1(LEU+)
2- Transformation
cdc13Δ stn1Δ ten1Δ
+ p-CDC13-TEN1-STN1 (URA+)
cdc13Δ stn1Δ ten1Δ
+ p-CDC13-TEN1-STN1 (URA+)
p-cdc13-ten1-stn1(LEU+)
cdc13Δ stn1Δ ten1Δ
p-cdc13-ten1-stn1(LEU+)
3- Growth on 5-FOA
Selection of ura- cells
4- Selection of ts mutants + plasmid recovery (yeast to bacteria)
+ transformation back into original strains
24°C
36°C
cdc13Δ stn1Δ ten1Δ
p-cdc13-ten1-stn1 ts (LEU+)
B
cdc13Δ stn1Δ ten1Δ
+ p-CST1 (URA+)
cdc13Δ stn1Δ ten1Δ
p-cst1-1 (LEU+)
cdc13Δ stn1Δ ten1Δ
p-cst1-4 (LEU+)
cdc13Δ stn1Δ ten1Δ
p-cst1-5 (LEU+)
cdc13Δ stn1Δ ten1Δ
p-cst1-15 (LEU+)
24°C
29°C
32°C
34°C
10 µm
34°C
C
cst1-15
Y388C
F684L
F865L
L11S
E32G
S105L
E225G
cst1-5
N180S;L190S
L392P;N405D
F832L
L11S
M320T
S364P
I412M
cst1-4
G273R
F510L;Y522C;D546G
L11S
S78P;M85T
L187S
cst1-1
S20P
N780S
L11S
CDC13
TEN1
STN1
1
924
1
160
1
494

### Figure S1

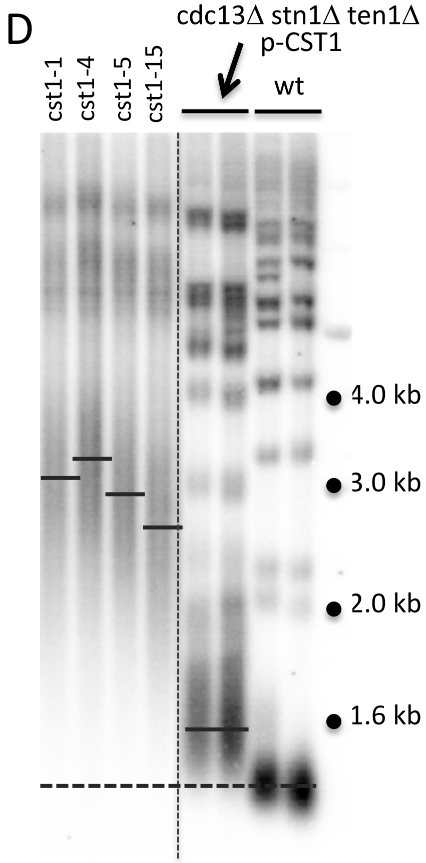


**Figure S1.** Construction and characterization of *cdc13Δ stn1Δ ten1Δ* mutants expressing temperature-sensitive *cdc13-ten1-stn1* fusion mutant genes. (**A**) A *cdc13Δ stn1Δ ten1Δ* triple deletion mutant of the CST complex surviving owing to the *CDC13-TEN1-STN1* synthetic *CST1* gene (fusion of wild-type genes; see Supplementary Materials and methods, section 1.1*)* expressed from a low-copy (centromeric) *URA3*$^{+}$ plasmid, YCplac33, was co-transformed with the PCR-mutagenized *cdc13-ten1-stn1* fusion and the gapped YCplac111-*CDC13-TEN1-STN1* (*LEU2*$^{+}$) fusion plasmid to allow gap repair using the mutagenized PCR. The YCplac33-*CDC13-TEN1-STN1* (*URA3*$^{+}$) wild type fusion plasmid was then shuffled out following growth on 5-FOA medium. The resulting *LEU2*$^{+}$ *ura*$^{-}$ *cdc13Δ stn1Δ ten1Δ* transformants, now relying only on mutagenized YCplac111-*cdc13-ten1-stn1* for survival, were then replica-plated onto YEPD plates both at 24 and 36°C and potential temperature-sensitive clones were isolated from the 24°C master plates, and their YCplac111-*cdc13-ten1-stn1* plasmid recovered by plasmid recovery (transfer from yeast to bacteria). Their temperature-sensitivity-conferring phenotype was re-checked following transformation back into the original *cdc13Δ stn1Δ ten1Δ* YCplac33-*CDC13-TEN1-STN1* (with again use of 5-FOA). These experiments were repeated several times, leading to the isolation of four mutants, as shown below. (**B**) The selected mutants of CST exhibited temperature-sensitive decreases in cell viability at temperatures comprised between 24 and 34°C, as assessed by performing "drop tests" (top panels), compared with the *cdc13Δ stn1Δ ten1Δ* mutant expressing the *CDC13-TEN1-STN1* hybrid gene (wild-type with no mutations), illustrated in leftmost top panels. In addition, at the restrictive temperature of 34°C, the *cdc13Δ stn1Δ ten1Δ p-cst1* mutants exhibited a typical G2/M arrest, as observed under the fluorescence microscope following DAPI staining (bottom panels, as indicated), *cdc13Δ stn1Δ ten1Δ* harboring wild-type YCp-*CDC13-TEN1-STN1* being illustrated under the same conditions (growth at 34°C) in leftmost bottom panels. Cell contours were outlined in order to better appreciate the position and stage of the nuclear material. (**C**) Sequences of the Cst1-1, Cst1-4, Cst1-5 and Cst1-15 mutant proteins. We note that all four alleles harbored the Ten1 L11S mutation and we verified that this mutation was never present in Ten1 of several wild-type strains. (**D**) Southern blot showing telomere organization and telomere length of various *cdc13Δ stn1Δ ten1Δ p-cst1* mutants, as indicated, compared with wild-type strains (wt) and with a *cdc13Δ stn1Δ ten1Δ* triple deletion mutant expressing the wild-type *CDC13-TEN1-STN1* fusion (*cdc13Δ stn1Δ ten1Δ p-CST1*). Telomere tracts of wt cells appear as a broad band of ~ 1.1-1.2 Kb (that includes the 300-350-nt-long terminal $TG_{1-3}$ telomere tracts). For each mutant, a horizontal bar has been drawn to indicate approximately the average size of the very large smear representing the average telomere length, for better comparison with the wt (horizontal dashed line). Teloblots shown here were from mutant cells grown at the permissive temperature for growth of 29°C.

genes would be very unlikely to complement a defect of a triple *cdc13*Δ *stn1*Δ *ten*1Δ deletion strain, thus giving chances to isolate genes involved in CST functions other than the CST genes themselves. Moreover, we postulated that simultaneously introducing mutations in the whole CST complex, that is to say mutagenizing a *CDC13-TEN1-STN1* synthetic fusion gene (see next two paragraphs), might reveal novel functions of the CST complex independent from its subunit-specific functions.

A *cdc13*Δ *stn1*Δ *ten1*Δ triple mutant expressing the wild-type *CDC13-TEN1-STN1* fusion gene from a centromeric plasmid (described in the **Supplementary Materials and methods**, section 1.1; **Fig. S1A**) was perfectly viable (highlighting functionality of the *CDC13-TEN1-STN1* synthetic gene, referred to as *CST1* gene), as shown on the drop test viability assays at temperatures comprised between 24 and 36°C (**Fig. S1B**; leftmost top panel). Moreover, at 34°C, this strain contained cells that were at various stages of the cell cycle (small buds correspond to S phase, large buds to G2 and M phases, no buds to G1 phase) and did not exhibit any apparent morphological defects (**Fig. S1B**; leftmost bottom panel). Note that we chose to name this synthetic gene *CST1* rather than *CTS1* (which would have been more logical given the order of the genes in the fusion) because *CTS1* is an already deposited gene name (encoding endochitinase) and, in addition, because *CST* is the name of the complex composed of Cdc13, Stn1 and Ten1 and the name cst1 for these mutants is therefore more descriptive of their functions.

The *cdc13*Δ *stn1*Δ *ten1*Δ *p-CDC13-TEN1-STN1* strain was then used to construct temperature-sensitive mutants, in which the wild-type *CDC13-TEN1-STN1* fusion gene was replaced by a PCR-mutagenized *cdc13-ten1-stn1* fusion introduced by gap repair (**Fig. S1A**), as also described in the **Supplementary Materials and methods**, section 1.1. We could isolate four temperature-sensitive mutants, so-called *cdc13*Δ *stn1*Δ *ten1*Δ *cst1* (referred to as *cst*Δ *p-cst1*), namely *cst*Δ *p-cst1-1*, *cst*Δ *p-cst1-4*, *cst*Δ *p-cst1-5* and *cst*Δ *p-cst1-15* that all exhibited growth defects at 32 or 34°C (**Fig. S1B**; top panels). All four mutants exhibited large-budded cells at the restrictive temperature of 34°C, typical of a G2/M arrest (**Fig. S1B**; bottom panels), in contrast with the *cdc13*Δ *stn1*Δ *ten1*Δ strain bearing the wild-type *CST1* gene, as seen above (**Fig. S1B**; leftmost bottom panel). All four *cdc13-ten1-stn1* fusion mutant alleles were sequenced (**Fig. S1C**). Of note, all four cst1 alleles harbored the L11S Ten1 mutation. We verified that this mutation was never present in the endogenous Ten1 of several wild-type strains. In addition, expressing (from a centromeric plasmid) either L11S *ten1* (the only *ten1* mutation in *cst1-15*) or the L11S, S78P and M85T *ten1* mutations present in *cst1-4* in a *ten1*Δ strain did not confer any apparent morphological or growth defects (**data not shown**). We also verified that the *CDC13-TEN1-STN1* fusion gene expressed from a centromeric plasmid perfectly rescued the temperature sensitivity of all four *cst*Δ *p-cst1* mutants. Telomere length measurement using the classical $^{32}$P-based telomere restriction fragment analysis method (TRF) established that all the isolated *cst*Δ *p-cst1* mutants exhibited strong deregulation of telomere size control (even at the permissive temperature for growth of 29°C), more precisely telomere lengthening to different degrees (**Fig. S1D**).

*2.2. Isolation of* SIZ1 *and* TOP2 *as extragenic suppressors of novel mutants of the telomeric Cdc13-Stn1-Ten1 complex*

In genetic screens in which genomic DNA librairies were transformed into the four temperature-sensitive *cst*Δ *p-cst1* mutants described above, *SIZ1* and *TOP2* were isolated as extragenic suppressors (see **Supplementary Materials and methods**, section 1.2; **Fig. S2A**). In the field of yeast genetics, suppression of growth defects by extragenic suppressors is more or less effective (meaning always partial) and can be assessed by performing growth assays, the so-called "drop tests" or "spot assays". A rescue phenotype such as observed here means improved growth capacity and increased survival in face of this telomeric damage upon overexpression of *SIZ1* or *TOP2. SIZ1* or *TOP2* overexpression allowed some rescue of the temperature-sensitive growth defect in the *cst*Δ *p-cst1-1* and *cst*Δ *p-cst1-15* mutants (**Fig. S2A**). In addition, *SIZ1* or *TOP2* overexpression also rescued *stn1*Δ *p-cst1-4*, *stn1*Δ *p-cst1-15*, *ten1*Δ *p-cst1-1*, *ten1*Δ *p-cst1-4*, *ten1*Δ *p-cst1-5* and *ten1*Δ *p-cst1-15* (**Fig. S2A** and **data not shown**).

Of note, the rescuing activity of *TOP2* towards the *cst*Δ *p-cst1-15* mutant provided by the pFL-44L library plasmid contained only part of *TOP2* sequences, being truncated for the first 1,571 nucleotides (total ORF is 4,287 nucleotides). Interestingly, this truncated *TOP2* gene rescued the *cst*Δ *p-cst1* mutants basically as efficiently as the full length *TOP2* gene cloned separately in a high-copy plasmid of the same family as the library plasmid (**Fig. S2B**).

Interestingly, *TEN1* was frequently found to rescue all four *cst*Δ *p-cst1* mutants (a total of 12 genomic fragments isolated; **Fig. S2B**), but *CDC13* and *STN1* were never isolated. Overexpression of *CDC13* or *STN1* alone from a multicopy plasmid similar to those from the genomic libraries did not rescue the *cst*Δ *p-cst1* mutants (**data not shown**), thus suggesting that the failure to isolate either one of these genes in the genetic screens was not due to the fact that they were simply missing from the libraries.

### Figure S2

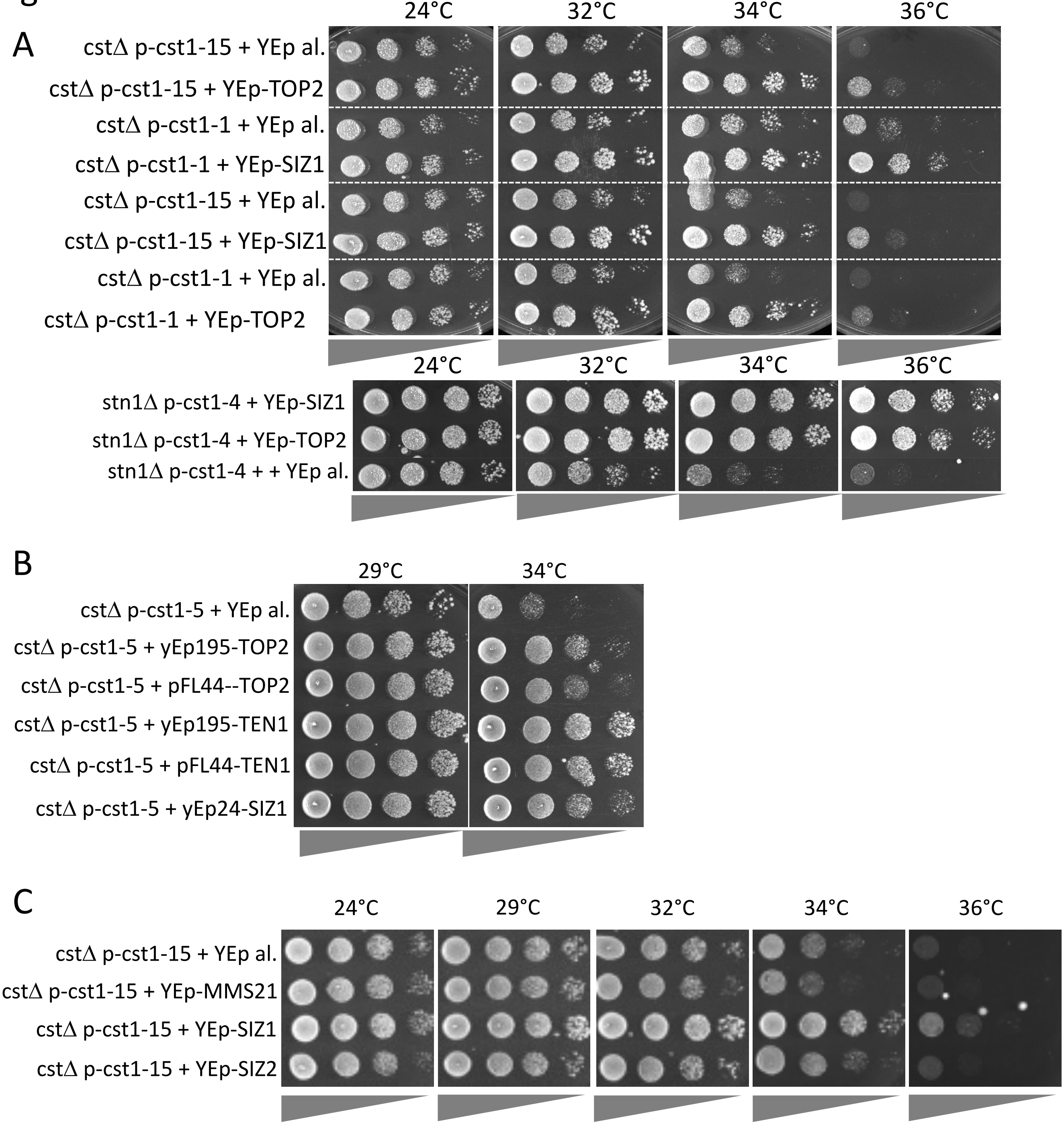


**Figure S2.** *SIZ1* and *TOP2* rescue mutants of the Cdc13-Stn1-Ten1 complex. (**A**) Overexpression of either *SIZ1* or *TOP2* from a 2μ multi-copy vector (YEplac195, labeled “YEp”) under their natural promoter and post-stop sequences resulted in improved growth of temperature-sensitive *cdc13*Δ *stn1*Δ *ten1*Δ triple deletion mutants harboring a temperature-sensitive *cdc13-ten1-stn1* fusion allele (*cst*Δ *p-cst1*, top panels) or of a *stn1*Δ null mutant expressing the temperature-sensitive *cst1-4* fusion allele (*stn1*Δ *p-cst1-4*, bottom panels). (**B**) Rescue of the *cst*Δ *p-cst1-5* mutant by overexpression of truncated *TOP2* (expressed from the pFL-44L bank plasmid) or of entire *TOP2* (recloned into YEplac195), or of *TEN1* (either from the original pFL-44L bank plasmid or recloned into YEplac195), or of *SIZ1* (from the YEp24 bank plasmid), together with negative control (vector alone, labeled “YEp al.”). (**C**) Overexpression of *MMS21*, coding for another SUMO E3 ligase, did not rescue at all the *cst1* mutants (in a *cdc13*Δ *stn1*Δ *ten1*Δ background), while rescue by *SIZ2* overexpression was well visible but much weaker than upon *SIZ1* overexpression.

Co-overexpressed *SIZ1* and *TOP2* did not rescue the defects of the *cst*Δ *p-cst1-1* and *stn1*Δ *p-cst1-4* mutant strains better than *SIZ1* or *TOP2* alone (**data not shown**). However, it should be noted that in some (**Fig. S2A**, top panels), but not all (**Fig. S2A**, bottom panels) of the tested mutants, overexpression of *SIZ1* rescued better than that of *TOP2*, thereby suggesting that Siz1 might act on an additional target other than Top2. This would also explain why overexpression of *TOP2* could not improve the rescue of the mutants when co-overexpresssed with *SIZ1*, Top2 being possibly already at its maximal level of sumoylation by Siz1 upon overexpression of *SIZ1* alone.

Interestingly, overexpression of *SIZ2* also rescued the *cst*Δ *p-cst1-15* mutant, but much less efficiently than overexpression of *SIZ1*, while overexpression of *MMS21* (coding for yet another SUMO E3 ligase) did not rescue at all that mutant (**Fig. S2C**). Rescue of the *cst*Δ *p-cst1* mutants by *SIZ2* was expected (assuming that sumoylation of Top2 is involved in the rescue mechanism; see Fig. 4 of main text), since both Siz1 and Siz2 can sumoylate Top2 [8]. The present finding that *SIZ1* was more efficient than *SIZ2* might suggest that Siz2 only acts on Top2 in the rescue mechanism, while Siz1 would have an additional target, most likely the septins, which are not sumoylated by Siz2 [5, 6]. However, it is equally possible that Siz1 provokes more sumoylation of Top2 than Siz2 does, which was shown previously to be the case [8].

In summary, the genetic experiments presented above established that both overproduction of Siz1 or Siz2, but not that of the third identified E3 SUMO ligase, Mms21, could rescue the temperature-sensitive defects of the *cst*Δ *p-cst1* mutants.

*2.3. Sumoylation of Top2 in the* stn1-sz2 *mutant and of the Cdc3 septin in the* ten1-sz, stn1-sz *and* cdc13-sz *mutants*

We were able to detect Myc2-Smt3 (Smt3 is *S. cerevisiae* SUMO) attached to Top2-HA$_2$, both tagged at their respective genomic locus, upon anti-HA immunoprecipitation, a *top2-SNM*-HA$_2$ Myc$_2$-*SMT3* strain serving as a control (**Fig. S3A**), in agreement with previous findings [9]. However, Top2 sumoylated bands were very similar in the *stn1-sz2* mutant and the wild type, thus leaving us unable to conclude that Top2 sumoylation is triggered or at least increased in response to CST loss of function.

Concerning septin sumoylation, first we set out to reproduce experimental conditions previously described in the literature concerning sumoylation of the Cdc3 septin in cells arrested in mitosis following nocodazole treatment ([4-6, 10]; see **Fig. S3B**). The *stn1-sz2* and *stn1*Δ *p-cst1-4* as well as the *ten1-sz* and *cdc13-sz* mutants exhibited highly increased levels of HA$_2$-Cdc3 sumoylation at the restrictive temperature of 34°C (**Fig. S3C**, third and fourth rows panels), just like the classical *cdc13-1* and *stn1-13* mutants (**Fig. S3C**, second row panels). Importantly, since septin sumoylation took place throughout mitosis in wild-type cells in the absence of any inflicted damage (**Fig. S4**; an event that we could establish here for the first time), it was not possible to determine whether HA$_2$-Cdc3 sumoylation occurring at 34°C in the mutants was due to their intrinsic damage or, rather, to the mitotic arrest or slowing down of these mutants. Noticeably, in these new mutants of CST, HA$_2$-Cdc3 sumoylation signals were also increased at 24°C (**Fig. S3C**, first, third and fourth rows panels), but, strikingly, were not increased in the classical *cdc13-1* and *stn1-13* mutants at 24°C (**Fig. S3C**, second row panels). As these were asynchronous cells, we next attempted to synchronize *stn1-sz2* and *cdc13-sz2* mutants with alpha factor in order to verify the presence or not of increased HA$_2$-Cdc3 sumoylation during cell cycle stages out of mitosis. However, even after 4 hr of treatment with alpha factor (normal treatment is 2 hr), the *stn1-sz2* and *cdc13-sz2* mutant cells failed to properly synchronize in G1 (Grandin and Charbonneau, unpublished data). Consequently, we strongly suspect that in asynchronous mutant cells at 24°C, there might be a much higher number of cells in mitosis than in wild-type asynchronous cells, hence the increase in septin sumoylation out of mitosis in these mutants.

#### 3.1 Discussion

*3.1. Possible mechanisms responsible for the rescue of the cst1 mutants by* SIZ1 *or* TOP2 *overexpression*

Two rather opposed hypotheses could be proposed to explain the rescue of the *cst1* mutants by overexpressed *SIZ1* or *TOP2*. First, Siz1 and Top2 might act to reinforce a particular cell cycle checkpoint activated by telomeric damage, thereby preventing excessive growth in the presence of telomeric damage, which gives them time to partially repair the damage to continue cell division, thus preserving cell viability and improving growth. This has been found to be the case for Mps1, a protein kinase that phosphorylates Mad1 and activates the spindle checkpoint. Overexpression of *MPS1* provoked an arrest of cells in mitosis in a Mad1-, Mad2-, Mad3-, Bub1-, Bub2- and Bub3-dependent manner even in the absence of spindle damage [11].

Alternatively, *SIZ1* or *TOP2* overexpression might provoke inactivation of a particular cell cycle checkpoint, thereby forcing the passage through the checkpoint-activated cell cycle arrest, resulting in improved growth, albeit with more intrinsic damage, but that might not obligatorily have

### Figure S3

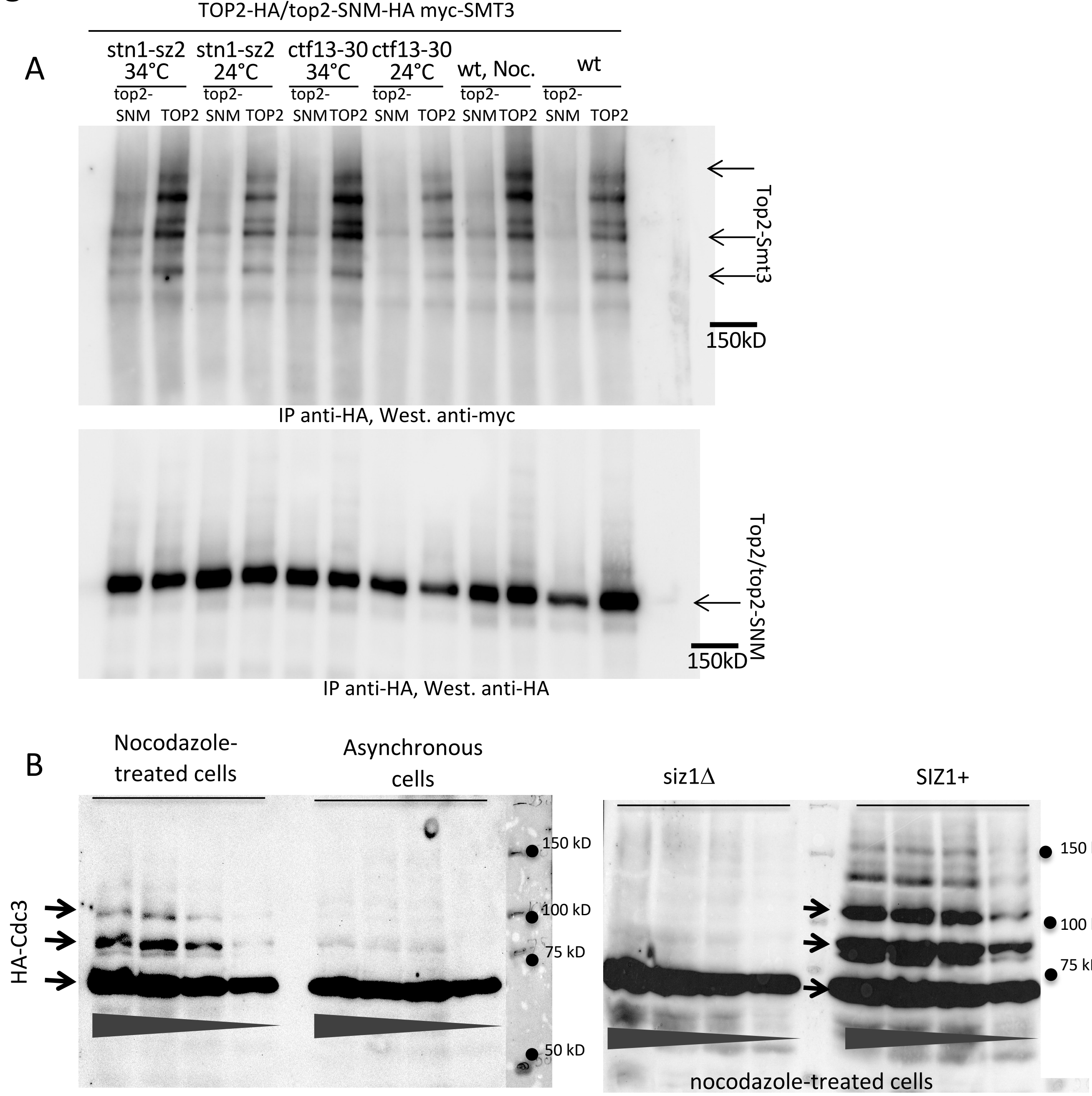


**Figure S3. (A)** Sumoylation of Top2-HA$_2$ in wild type cells (wt) treated or not with 15 μg/ml nocodazole (Noc.) as well as in the temperature-sensitive *stn1-sz2* and *ctf13-30* mutants, under the indicated conditions. All these strains had their genomic *TOP2* locus tagged with HA, as well as their genomic *SMT3* (SUMO) locus tagged with Myc. In parallel, the same strains harboring *top2-SNM* mutation at their genomic *TOP2* locus served as a control to identify the bands representing sumoylated Top2 protein. See also Supplementary Materials and methods, section 1.3). (**B**) Septin sumoylation was measured in asynchronous wild-type cells expressing HA$_2$-*CDC3* from *CDC3* genomic locus. For each of the four panels, decreasing amounts of the same extract were loaded from left to right (as indicated by the bars at the bottom; same dilution factor for extracts of the same experiment, left or right panel). Asynchronous untreated cells (right part of left panel) exhibited no signal of sumoylation, just like *SIZ1*-deleted cells (*siz1*Δ) treated with 15 μg/ml nocodazole for 2 hr (left part of right panel), while treatment with nocodazole in the *SIZ1*$^+$ background (left part of left panel and right part of right panel) arrested cells in metaphase of mitosis, which triggered massive HA$_2$-Cdc3 sumoylation. Bottom arrows represent the HA-tagged non-sumoylated form of Cdc3, while the top two arrows represent the two major HA$_2$-Cdc3 sumoylated forms, observed in all experiments. In the right part of right panel, the two upper bands (sumoylated, because absent in the *siz1*Δ control at left) were not detected in all experiments, probably because of a low dilution of the extract used here.

### Figure S3

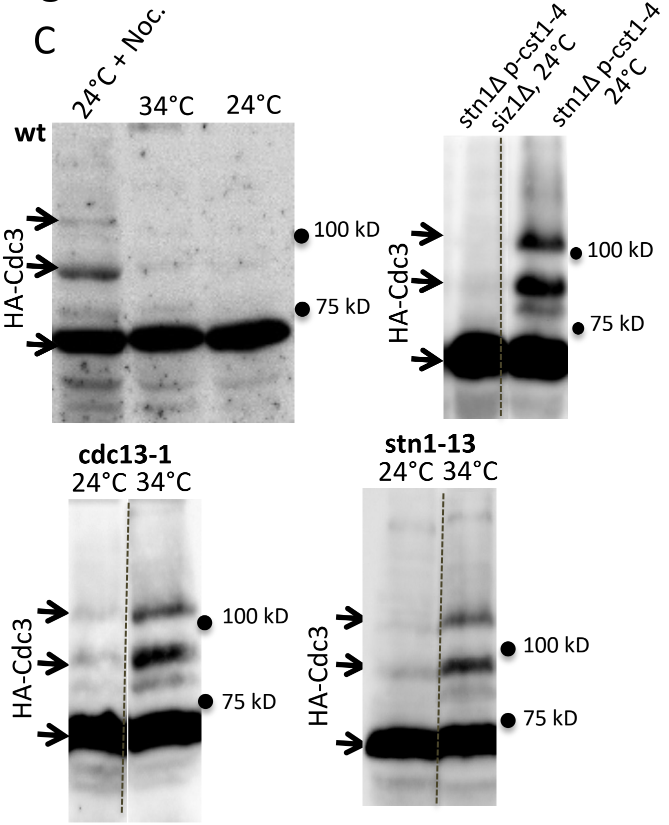


**Figure S3.** (**C**) $HA_2$-Cdc3 sumoylation signals (two major forms of sumoylation indicated by top two arrows) in the wild type (wt) and mutants of CST, as indicated and under the indicated conditions. Importantly, measurement of $HA_2$-Cdc3 sumoylation in the *stn1*Δ *p-cst1-4 siz1*Δ double mutant (first row panels) allowed determination of the sumoylated or non-sumoylated nature of the $HA_2$-Cdc3 bands.

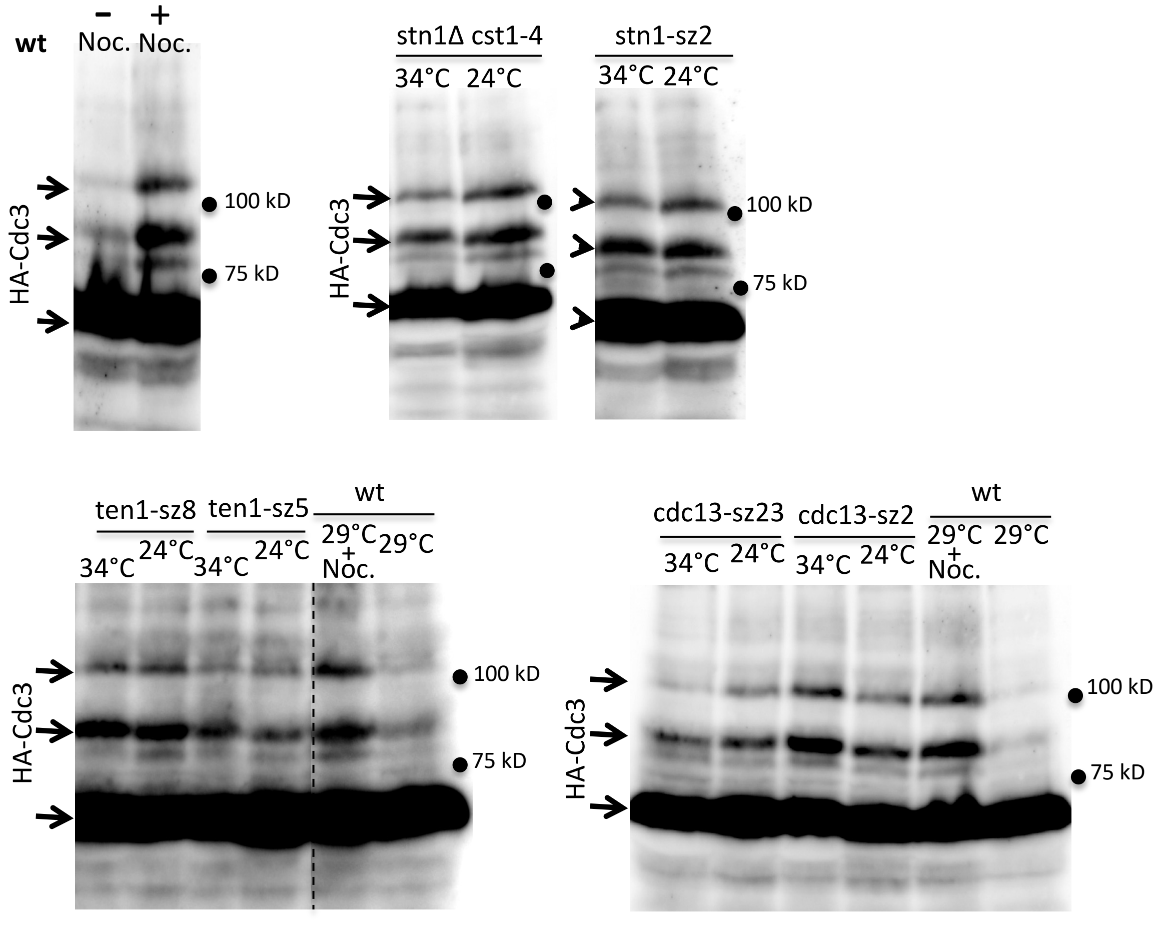

Figure S4

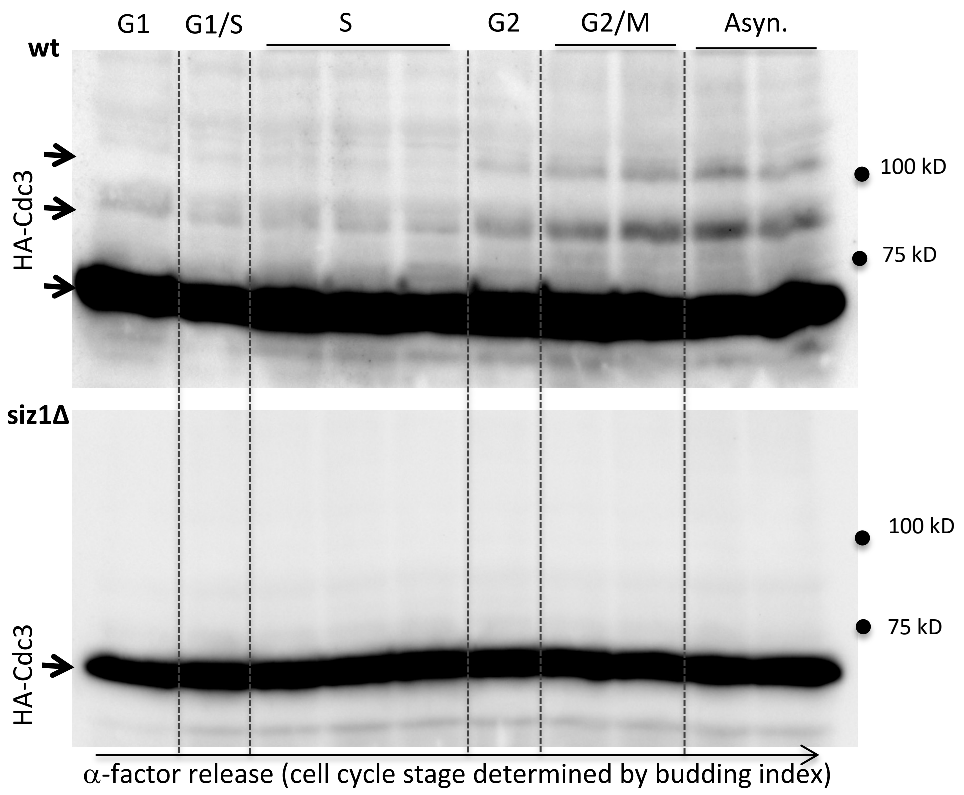


**Figure S4.** In wild-type (wt) alpha-factor-synchronized cells, $HA_2$-Cdc3 sumoylation peaked at G2/M and stayed very low during the rest of the cell cycle. In **top panel**, the bottom arrow represents the $HA_2$-tagged non-sumoylated form of Cdc3, while the top two arrows represent the two major $HA_2$-Cdc3 sumoylated forms. In **bottom panel**, measurements of $HA_2$-Cdc3 sumoylation in a *SIZ1*-deleted strain (*siz1*Δ) confirm the sumoylated nature of the $HA_2$-Cdc3 bands present in the wt, above. The arrow indicates the position of the $HA_2$-tagged non-sumoylated form of Cdc3, as above.

dramatic consequences in the short term. Such mechanisms have already been described concerning the *cdc13-1* telomeric mutant, which, at semi-permissive temperatures for growth, forms microcolonies when the DNA damage checkpoint has been genetically inactivated [12]. Similarly, overproduced Cdc20 (APC-Cdc20 is the target of the Mad2 spindle checkpoint) was seen to override the Rad9-mediated DNA damage checkpoint in the *cdc13-1* mutant [13]. In addition, *cdc13-1* arrest could be rescued when *SIC1*, coding for an inhibitor of Cdc28-Clb2, was overexpressed [14]. Moreover, overproduction of Cdh1, a member of the APC-Cdh1, under the control of the strong, constitutive *GAL1* promoter, could force escape from a nocodazole arrest or an arrest provoked by septin inactivation [15, 16]. Finally, in humans, it was found that overexpression of AURORA-A might interfere with spindle-microtubule attachment and disrupt regulation of the spindle checkpoint by allowing cells with abnormal chromosomal separation to enter anaphase [17].

Additional types of mechanisms could be invoked. For instance, overexpressed *TOP2* was proposed to be mainly produced under its non-sumoylated form and to titrate out the experimentally produced sumoylated forms of Top2 in the *cdc13-1 smt4*Δ mutant (official name for *SMT4* is now *ULP2*), thereby rescuing the *smt4*Δ-induced cohesion defect, Smt4 being a SUMO isopeptidase that reverses the levels of Top2 sumoylation [9]. In a distinct study, overexpressed *TOP2* could rescue a null mutation of *PDS5* (coding for a component of the cohesin complex), but, this time, independent of the *pds5*Δ-induced cohesion defect [18]. In that situation, Top2 might nevertheless cooperate with Pds5, as deduced from the analysis of the *pds5*Δ *top2-4* double mutant, to promote proper chromosome segregation, overproduced Top2 providing increased decatanation of sister chromatids compensating for diminution of sister chromatid separation provoked by *pds5*Δ [18]. Therefore, in the present study, overexpressed *TOP2* might also act by titration of hyper-sumoylated forms of Top2 provoked by the *cst1* mutations. Alternatively, overexpressed *TOP2* might directly act upon molecular events parallel and complementary to events rendered dysfunctional by the *cst1* mutations, thus reinforcing a common target.
